# Comparison of phage-derived recombinases for genetic manipulation of *Pseudomonas* species

**DOI:** 10.1101/2023.05.16.541022

**Authors:** Madison J. Kalb, Andrew W. Grenfell, Abhiney Jain, Jane Fenske-Newbart, Jeffrey A. Gralnick

## Abstract

Several strains in the *Pseudomonas* genus are categorized as plant growth promoting rhizobacteria (PGPR). Although several of these strains are strong candidates for applications as biofertilizers or biopesticides, known genome editing approaches are generally limited and require further development. Editing genomes in PGPR could enable more robust agricultural applications, persistence and biosafety measures. In this study, we investigate the use of five phage-encoded recombinases to develop a recombineering workflow in 3 PGPR strains: *P. protegens* Pf-5, *P. protegens* CHA0, and *P. putida* KT2440. Using point mutations in the *rpoB* gene, we reach maximum recombineering efficiencies of 1.5 x 10^-4^, 3 x 10^-4^, and 5 x 10^-5^, respectively, in these strains using λ-Red Beta recombinase from *E. coli*. We further examine recombineering efficiencies across these strains as a function of selected mutation, editing template concentration, and phosphorothiolate bond protection. This work validates the use of these tools across several environmentally and biotechnologically relevant strains to expand the possibilities of genetic manipulation in the *Pseudomonas* genus.

**Importance:** The *Pseudomonas* genus contains many members currently being investigated for applications in biodegradation, biopesticides, biocontrol and synthetic biology. Though several strains have been identified with beneficial properties, in situ genetic manipulations to further improve these strains for commercial applications have been limited due to lack of efficient genetic tools that have been tested across this genus. Here we test the recombineering efficiencies of 5 phage-derived recombinases across 3 biotechnologically relevant *Pseudomonas* strains: *P. putida* KT2440, *P. protegens* Pf-5, and *P. protegens* CHA0. These results demonstrate a method to generate targeted mutations quickly and efficiently across these strains, ideally introducing a method that can be implemented across the *Pseudomonas* genus and a strategy that may be applied to develop analogous systems in other non-model bacteria.

## Introduction

Much of our power to study and understand microorganisms lies in our ability to genetically modify them. The advent of genetic modification completely changed the field of microbiology: from enabling researchers to elucidate gene function and determine the role of genetic elements, to engineering new microorganisms for biotechnological applications. A relatively new method of genetic modification that has become increasingly popular due to its high fidelity and potential for multiplexing is recombination-mediated genetic engineering or recombineering.

Recombineering is a genetic modification method which utilizes prophage-derived single strand DNA annealing proteins (SSAPs) termed recombinases to introduce precise mutations into actively growing cells (1–3). This gene modification method revolutionized the field of genetics as it allowed scarless mutations to be introduced into either a plasmid or chromosome at a relatively low cost and with mutation rates comparable to other methods. The phage derived SSAPs form an oligomeric ring around single stranded DNA (ssDNA) and facilitate annealing to homologous DNA as an Okazaki fragment, requiring only 40-50 nucleotides of homology to the genetic target (2, 4–7). Recombineering can utilize dsDNA or ssDNA substrate, although dsDNA requires the addition of the SSAP’s complimentary exonuclease (2, 4). ssDNA recombineering is often preferred as oligonucleotide substrate can be customized and synthesized for relatively low cost, while also only requiring the expression of the SSAP.

The most widely studied SSAPs are λ-Red Beta from the *Escherichia coli* λ phage and RecT from the *E. coli* Rac prophage, though the functionality of these recombinases are fairly limited to closely related genera (8–11). Attempts at recombineering in other genera including *Lactobacillus, Corynebacterium, Pseudomonas* and even wild *E. coli* strains have not yet reached high levels of efficiency compared to model strains of *E. coli* (12–16). Difficulties with applying this platform in non-model organisms have been attributed in part to using hosts with less understanding and prior genome modifications than laboratory strains of *E. coli* as well as recombinase portability issues (9, 17).

Recombineering attempts in non-model organisms have been improved by the screening of λ-Red Beta and RecT homologs (5–7). Recent phylogenetic studies have identified six families of recombinases, though so far most efficient recombinases tested within recombineering frameworks are from the Rad52 superfamily, which includes both the λ-Red Beta and RecT proteins (7, 18, 19). Though recombineering efficiencies are impacted by many variables, an area of major focus is the recombinase, as this choice can influence efficiency by several orders of magnitude (7,8). Additional strategies to improve recombineering efficiencies are centered on oligonucleotide design, including optimizing homology arm length, eliminating hairpins or other secondary structures, minimizing off-target binding, and targeting to the replication fork lagging strand (2, 20).

In this work we aimed to develop a recombineering system in a selection of environmentally relevant *Pseudomonas* strains: *P. protegens* Pf-5, *P. protegens* CHA0, and *P. putida* KT2440, as these organisms can improve crop integrity and yields (21–25). Our strategy included screening several SSAPs for activity across these strains, as previous research for recombineering *Pseudomonas spp.* has primarily focused on a few strains and a few recombinases. Here we test a RecT homolog from the *P. syringae pv. syringae* B728a strain, which has been shown to facilitate recombineering in other Pseudomonads (26, 29), as well as four additional recombinases. We also investigate the effects of oligonucleotide amounts and mutation design on recombineering efficiency with the overall development of a recombineering system in these environmental *Pseudomonas* isolates.

## Materials and methods

### Bacterial strains and cultivation

All strains and plasmids used in this work are listed in Table 1. Lysogeny broth (LB) (BD Difco™ Dehydrated Culture Media: LB Broth, Miller) was used to routinely culture bacteria. When necessary, growth media was supplemented with kanamycin at a final concentration of 50 µg/mL for *P. protegens* Pf-5, *P. protegens* CHA0, and *E. coli*, while a concentration of 100 µg/mL was used for *P. putida* KT2440, or rifampicin at a concentration of 50 µg/mL for all strains. Media containing rifampicin was wrapped in foil to prevent photodegradation. Cultures were grown aerobically at 30°C (*Pseudomonas spp.*) or 37°C (*E. coli*) and shaken at 250 rpm when grown in liquid culture.

**Table 1:**
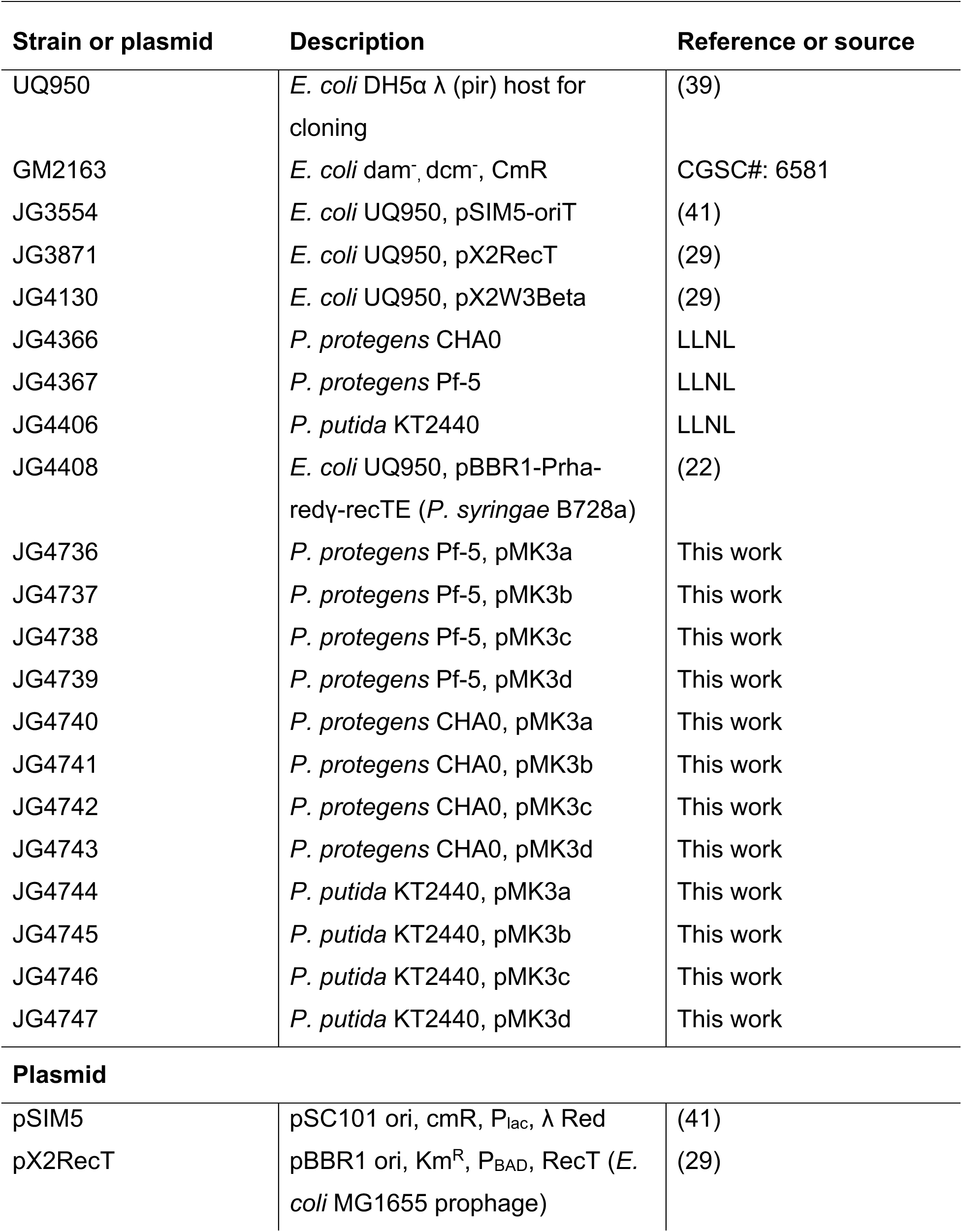
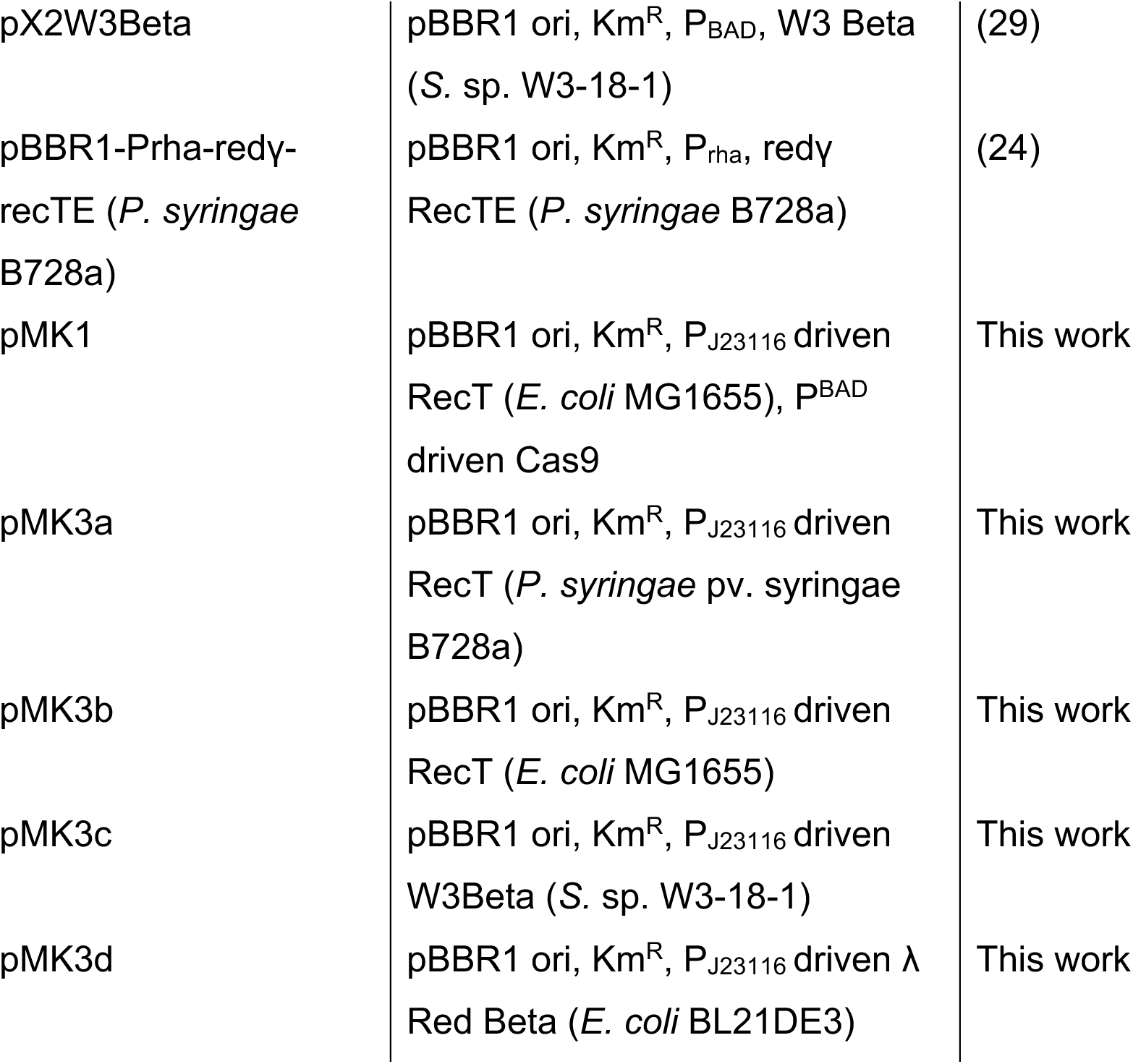
Strains and plasmids used in this work.

### Plasmid construction

Relevant sequences for plasmid construction are listed in Table 2. All primers used to construct plasmids are listed in Table 3. Primers were obtained from IDT (Coralville, IA). Cloning fragments were PCR amplified using Q5 polymerase 2X master mix (New England Biolabs). Full construct sequencing was performed by Plasmidsaurus (Eugene, OR). pMK1 was generated by Gibson assembly of the pX2Cas9 backbone with *E. coli* MG1655 RecT and a Gblock containing the T24 terminator sequence, P_J23116_ constitutive promoter, and RBS Sp17 from (28). *araC*, P^BAD^, and Cas9 were removed from pMK1 to generate pMK2 using PCR introduced BsaI sites. Recombinase genes were PCR amplified from JG3554 (*E. coli* BL21DE3 λ Red), JG3871 (*E. coli* MG1655 RecT), JG4130 (*S.* sp. W3-18-1), JG4408 (*P. syringae* B728a RecT) with flanking BsaI sites for Golden Gate Cloning into the pMK2 backbone downstream of RBS Sp17 to generate pMK3x plasmids to specifically test constitutive recombinase expression.

**Table 2.**
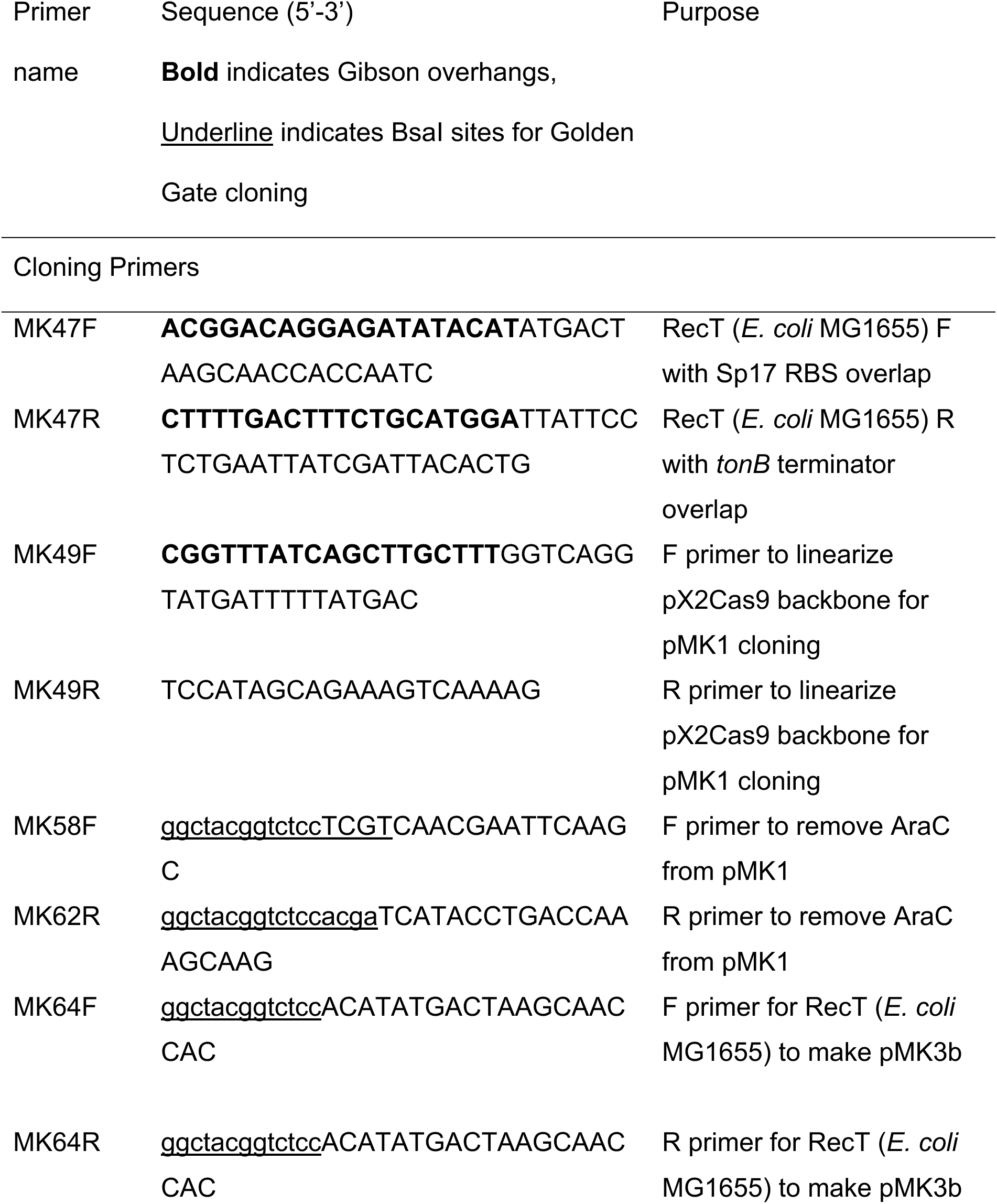
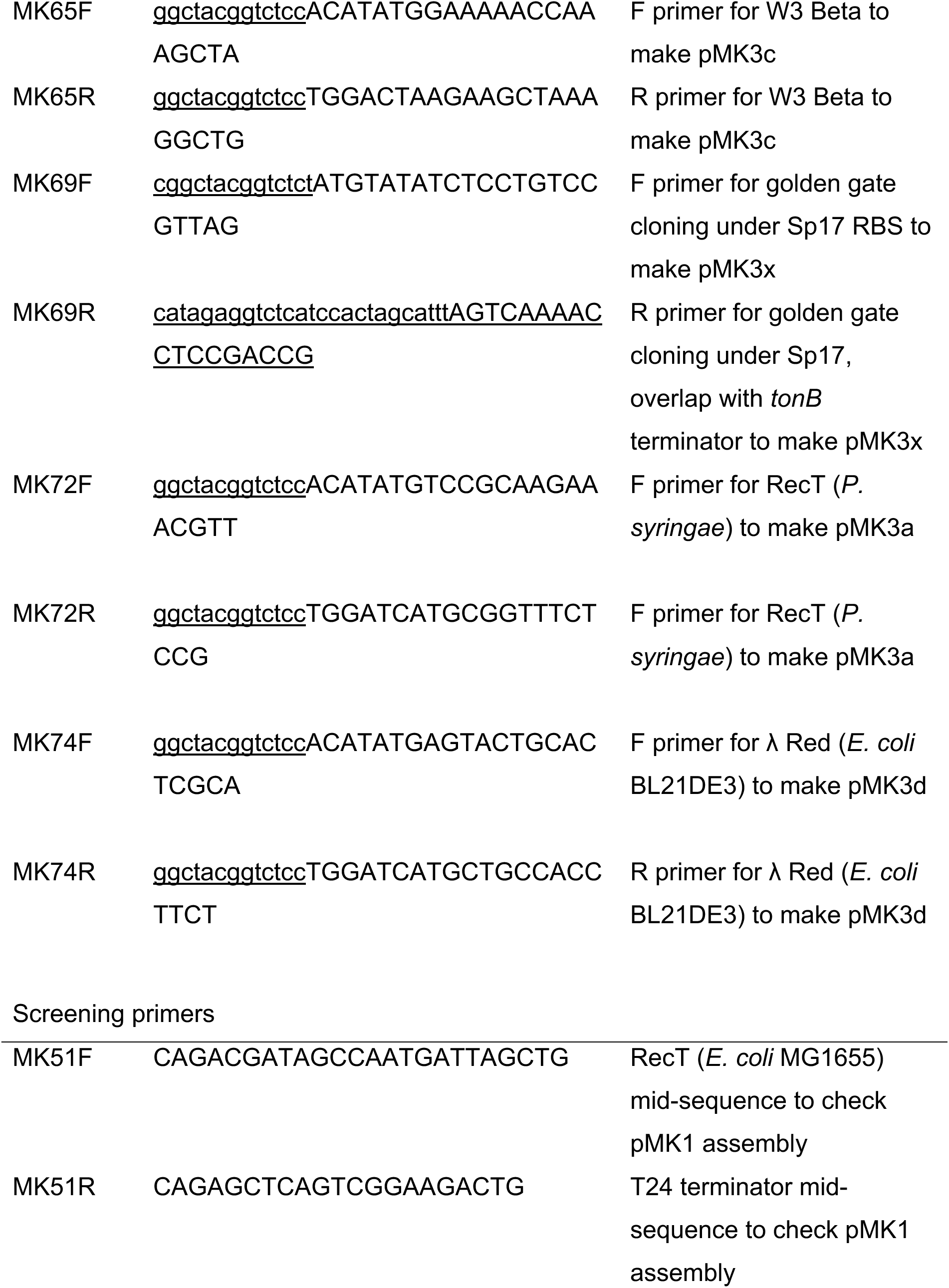
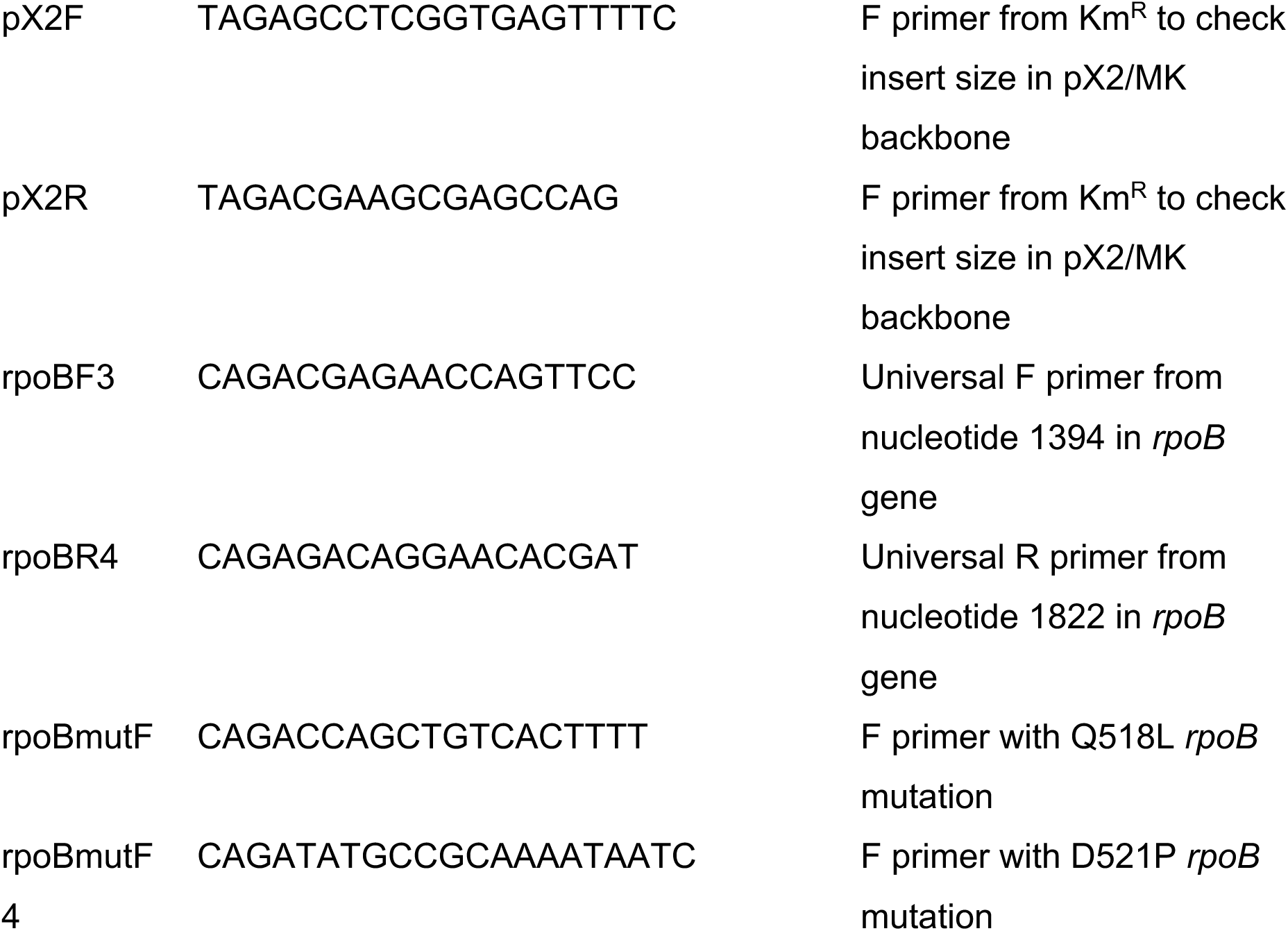
Primers used in this study.

**Table 3.**
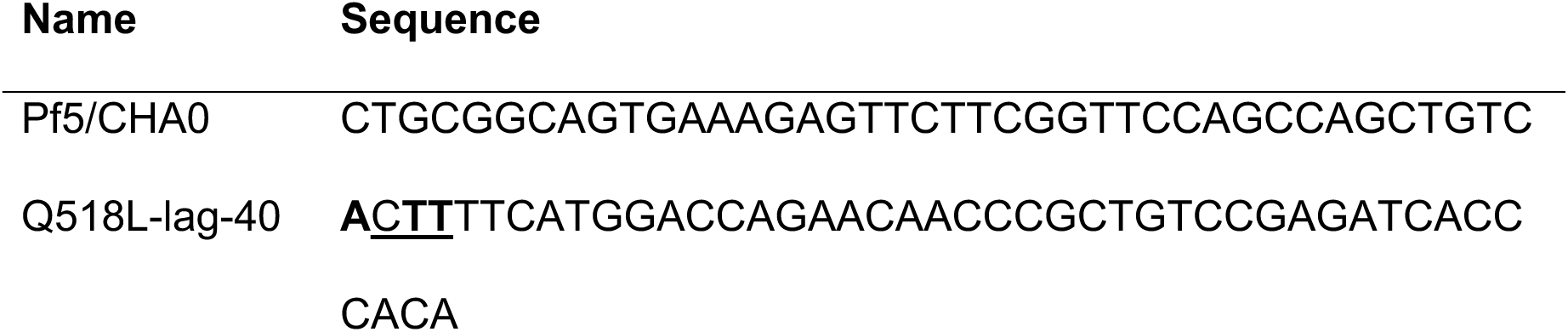
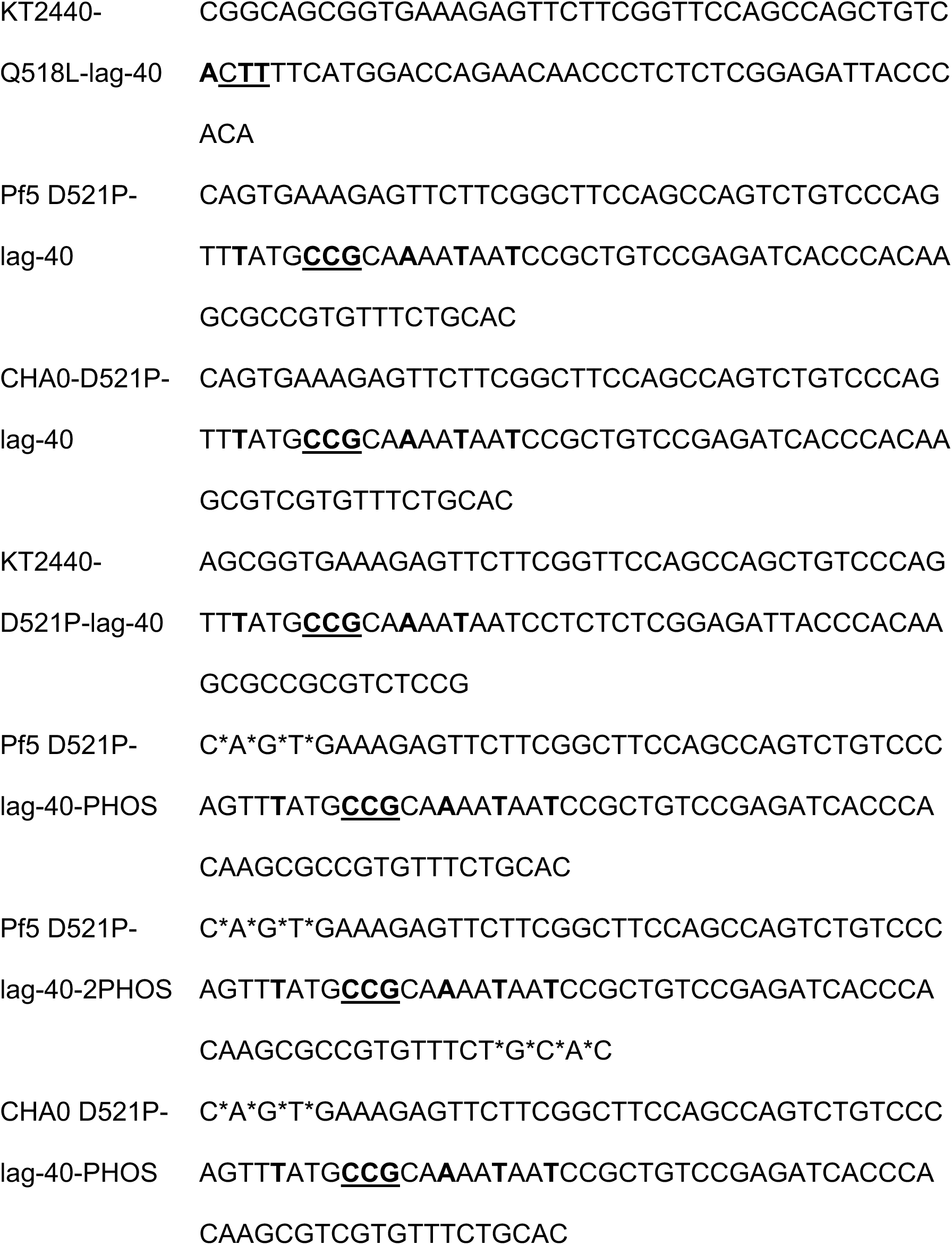
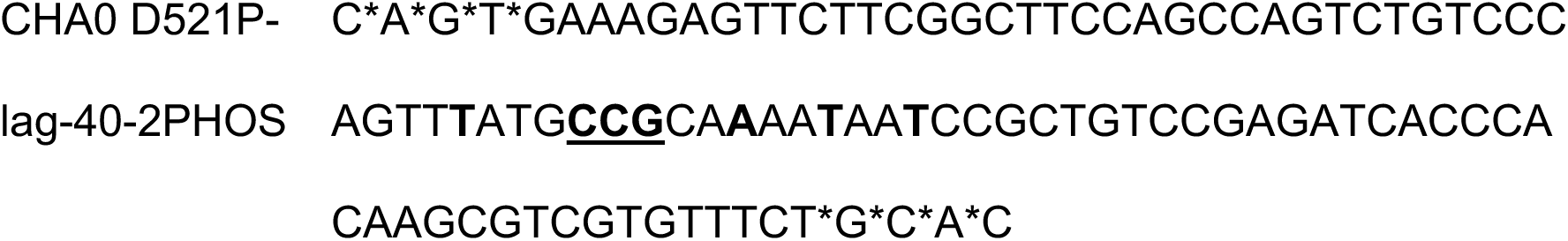
ssDNA oligonucleotides used for chromosomal recombineering. **Bold** indicates nucleotide mismatches to the wildtype sequence; underline indicates the codon carrying the specified point mutation; * indicates phosphorothioate bond.

### Plasmid transformation

For routine plasmid transformation, single colonies of the wildtype *Pseudomonas* strains were cultivated overnight at 30°C and 250 rpm. 2 mLs of overnight culture were harvested by centrifugation at 10,000 rpm for 1 minute, and electrocompetent cells were produced by gentle washing 3 times using 1 mL of either 1 M sorbitol (pH 7.6) for *P. protegens* Pf-5 and CHA0 or 300 mM sucrose for *P. putida* KT2440, as these electroporation buffers resulted in highest transformation efficiencies in these strains (Supplementary Figure S1). A minimum of 50 ng of plasmid DNA was introduced to the final resuspended volume of ∼60-70 µL electrocompetent cells, and this mixture was transferred to a 0.1 cm electroporation cuvette and a pulse was applied (settings: 25 μF; 200 Ω; 1.2 kV using the Bio-Rad GenePulserXcell™; Bio-Rad). 1 mL of LB was added, and cells were transferred to a 2 mL Eppendorf tube to recover at 30°C and 250 rpm for 2 hours. Cultures were then plated on selective media to isolate transformants and incubated at 28°C.

### Screening for recombineering target

To identify rifampicin-resistance (Rif^R^) mutations in our group of Pseudomonads, overnight cultures of each strain were plated on 50 μg/mL or 100 μg/mL rifampicin and incubated at 30°C until colonies formed (36-48 hours). Primers were designed to amplify clusters I and II of the *rpoB* gene (24), where rifampicin is known to bind and most Rif^R^ mutations occur. This 400 bp fragment within the *rpoB* gene of 10 Rif^R^ mutants for each strain grown on 50 µg/mL rifampicin was PCR amplified and sequenced to identify point mutations resulting in rifampicin resistance. Sequence outputs were aligned using the EMBL-EBI MUSCLE (MUltiple Sequence Comparison by Log-Expectation) tool.

### Oligonucleotide design

Recombinogenic oligonucleotides can be found in Table 3. Oligonucleotides were obtained as 250 nmole oligos from IDT (Coralville, IA) and resuspended in water to a final concentration of 100 µM. The oligonucleotides were designed to introduce single point mutations within the *rpoB* gene resulting in rifampicin resistance. Point mutations were flanked by silent mutations to evade methyl-directed mismatch repair (MMR). Guidelines for oligonucleotide design have been described elsewhere (15, 30). 40 bp of homology flanked each side of the mutagenic segment to generate recombinogenic oligonucleotides of 90-100 nucleotides in length. The mFold application via UNAFold (http://www.unafold.org/mfold/applications/dna-folding-form.php) was used to calculate DNA folding energies using a folding temperature of 30°C and the default settings (31, 32). Optimal folding temperatures of recombinogenic oligonucleotides in *E. coli* are above -20 kcal/mol, with a peak at about -12.5 kcal/mol (30). In this work a range of -7 to -14 kcal/mol was used. When indicated, four phosphorothiolate bonds were introduced at the 5’ end or both 5’ and 3’ ends of the oligonucleotide to investigate the effect of DNA protection against exonucleases.

### Recombinase comparisons in *Pseudomonas*

Plasmids pMK3a, b, c, d, or e were transformed into *P. putida* KT2440, *P. protegens* Pf-5, or *P. protegens* CHA0 via electrotransformation as indicated above. Single colonies were inoculated into LB and grown overnight at 30°C and 250 rpm. Cultures were diluted to an OD600 ∼ 0.085 and grown in the same conditions until they reached an OD of 0.4-0.7. 4 mLs of culture were used per replicate. Cells were made electrocompetent as above, and then 5 µL of 100 µM recombinogenic oligo carrying either the Q518L or D521P *rpoB* point mutations were mixed into the cell suspension. To improve recombineering likelihood, recovery time was extended to 3 hours, after which tenfold dilutions were plated onto LB rifampicin (50 µg/mL) as well as LB and incubated at 28°C. Colonies were counted after 2 days of growth except in the case of *P. protegens* Pf-5 and *P. protegens* CHA0 when the Q518L oligo was used, in which colonies were counted after 4 days of incubation. All experiments included at least 3 biological replicates. Presence of mutation was confirmed via mutation specific PCR primers rpoBmutF, rpoBmutF3, and rpoBmutF4 and reverse universal primer rpoBR4. Mutations were also confirmed using Sanger sequencing as discussed above from PCR reactions using primers rpoBF3 and rpoBR4.

### Optimization of recombineering

To further improve recombineering efficiencies, we investigated the effects of oligo availability impacted by oligo load or phosphorothioate bonds using the D521P point mutation and pMK3d plasmid. Cells were made electrocompetent as indicated above, and varying oligo amounts of 0.3, 3, 15, and 30 µg to test oligo load or 15 µg of oligo with four phosphorothioate bonds on either the 5’ end or both the 5’ and 3’ ends were introduced to the cell suspensions. Cells were electroporated, recovered, and plated as described above.

### Statistical and sequence analysis

Pairwise comparisons to calculate significance levels between groups means was performed using the Mann-Whitney U test. Multiple sequence alignments were made using Clustal Omega from EMBL-EBI using standard parameters.

## Results

### Identification of positive selection point mutations in *rpoB*

Our experimental setup was designed assuming a low frequency of recombinase-mediated allele integration by surveying a genetic target to select for recombinants against nonedited members within the population (Figure 1). Several positive selection targets have been explored in both *E. coli* and *Pseudomonas* spp., including the *rpsL, pyrF, tolC, gyrA,* and *rpoB* genes (7, 17, 28). In each of these targets, the incorporation of ssDNA encoding selective point mutations or premature stop codons can result in a genotype that can be selected for. This experimental setup can be used to determine recombineering efficiencies between candidate SSAPs as well as determine optimal recombineering conditions within each targeted strain.

**Figure 1.**
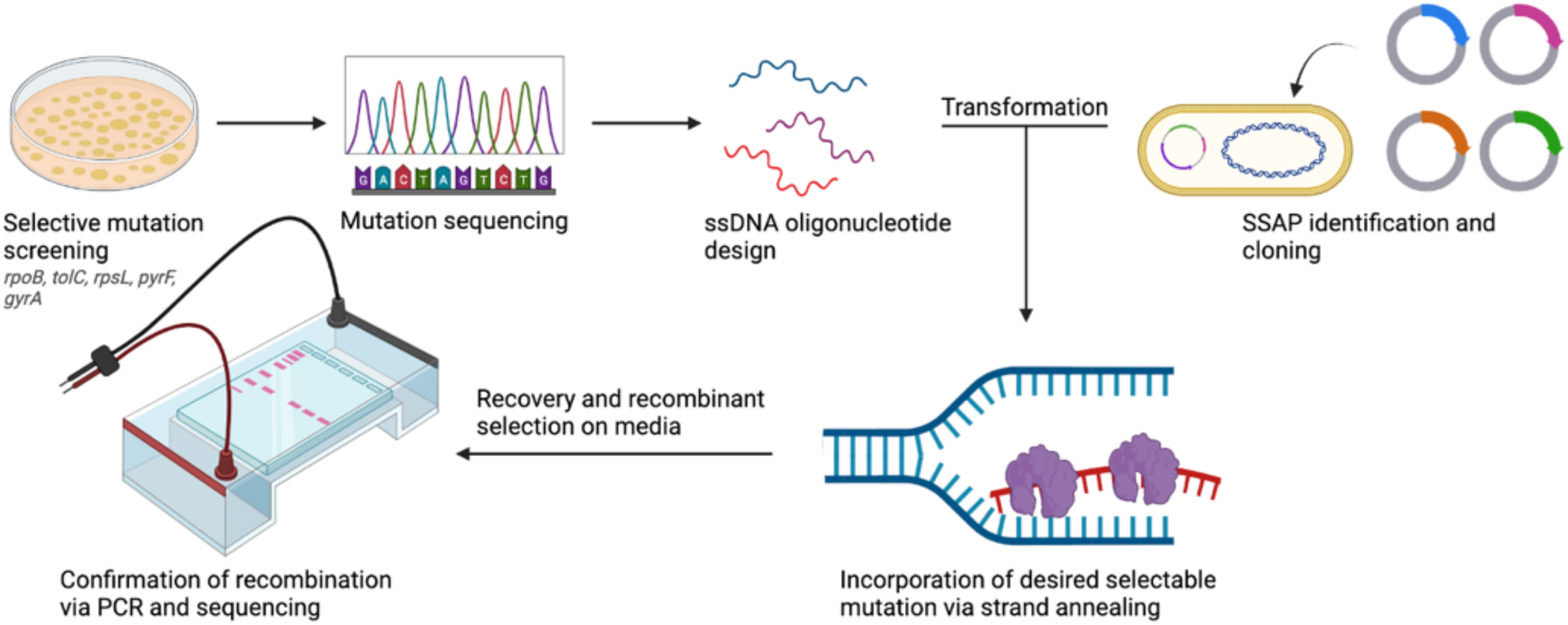
Overview of experimental setup. To determine recombineering efficiencies across different strains and conditions, we screened our *Pseudomonas* strains for rifampicin-resistance (Rif^R^) encoding mutations by sequencing Clusters I and II of *rpoB*. We then designed ssDNA oligonucleotides encoding our screened mutations with 40-basepair homology arms and introduced them into log-phase cultures expressing individual SSAP candidates. Efficiency of recombineering was calculated by normalizing number of Rif^R^ colonies to number of viable cells after recovery. Confirmation of intended mutation was performed using PCR and Sanger sequencing. Figure generated using BioRender.

We surveyed the *rpoB* gene to find rifampicin resistance encoding point mutations within our selection of *Pseudomonas* species. After 36-48 hours of growth, we sequenced 25 Rif^R^ colonies (10 from *P. protegens* Pf5, 5 from *P. protegens* CHA0, and 10 from *P. putida* KT2440) over Clusters I (amino acid 510-542) and II (amino acids 562-575) of the *rpoB* gene, as most Rif^R^ mutations are made in this region (29) (Table S1). All of the point mutations identified in this study were located in Cluster I, specifically between amino acids 517 and 536. A mutation at amino acid residue 521 occurred in all the strains tested, with the most common being an A to G transition mutation resulting in the exchange of glycine for aspartic acid. As this seemed a robust Rif^R^ mutation across all strains, we designed recombinogenic oligos to target this residue, as well as a previously reported Rif^R^ mutational residue at position 518 (29). A model of the RNA polymerase beta subunit binding pocket with both point mutations can be found in Supplementary Figure S2.

### Recombinase efficiency varies across *Pseudomonas* strains

Using the *rpoB* gene target, we tested a selection of five phage-derived recombinases in their ability to introduce a point mutation conferring rifampicin resistance in three strains: *P. protegens* Pf-5, *P. protegens* CHA0, and *P. putida* KT2440. The selected recombinases span a variety of SSAPs reported to function in different Gammaproteobacteria, all within the broader RecT family of recombinases (7, 26, 27, 33, 34). As the RecT family of recombinases was the most enriched under selective pressure in a previous study comparing SSAPs from all 6 major families, we selected candidates within that family to survey within our *Pseudomonas* strains (7). A few of these candidates had been tested to limited extent in other strains of *Pseudomonas*, however an investigation directly comparing multiple candidate SSAPs in this selection of strains had not yet been performed. A summary of select recombineering efforts to date in *Pseudomonas* can be found in Table S2.

To determine the relative efficiencies of each recombinase in the different *Pseudomonas* strains, we introduced an oligonucleotide encoding a D521P mutation in the *rpoB* gene in strain backgrounds harboring plasmids with constitutively expressed recombinase. Presence of the D521P mutation was determined using PCR amplification and sequencing of the *rpoB* gene region. Experiments using an empty vector strain and recombinase-carrying strains without oligonucleotides were used to determine background allelic exchange frequencies and spontaneous Rif^R^ frequencies, respectively. Spontaneous Rif^R^ frequencies for the three strains in the presence or absence of recombinase were 1.3 x 10^-7^ for *P. protegens* Pf-5, 5.8 x 10^-8^ for *P. protegens* CHA0, and 3.6 x 10^-7^ for *P. putida* KT2440 (Figure 2). In the absence of recombinase, allelic exchange frequencies with the addition of oligonucleotide ranged from 5 x 10^-7^ in *P. protegens* CHA0 to 8 x 10^-8^ in *P. putida* KT2440.

**Figure 2.**
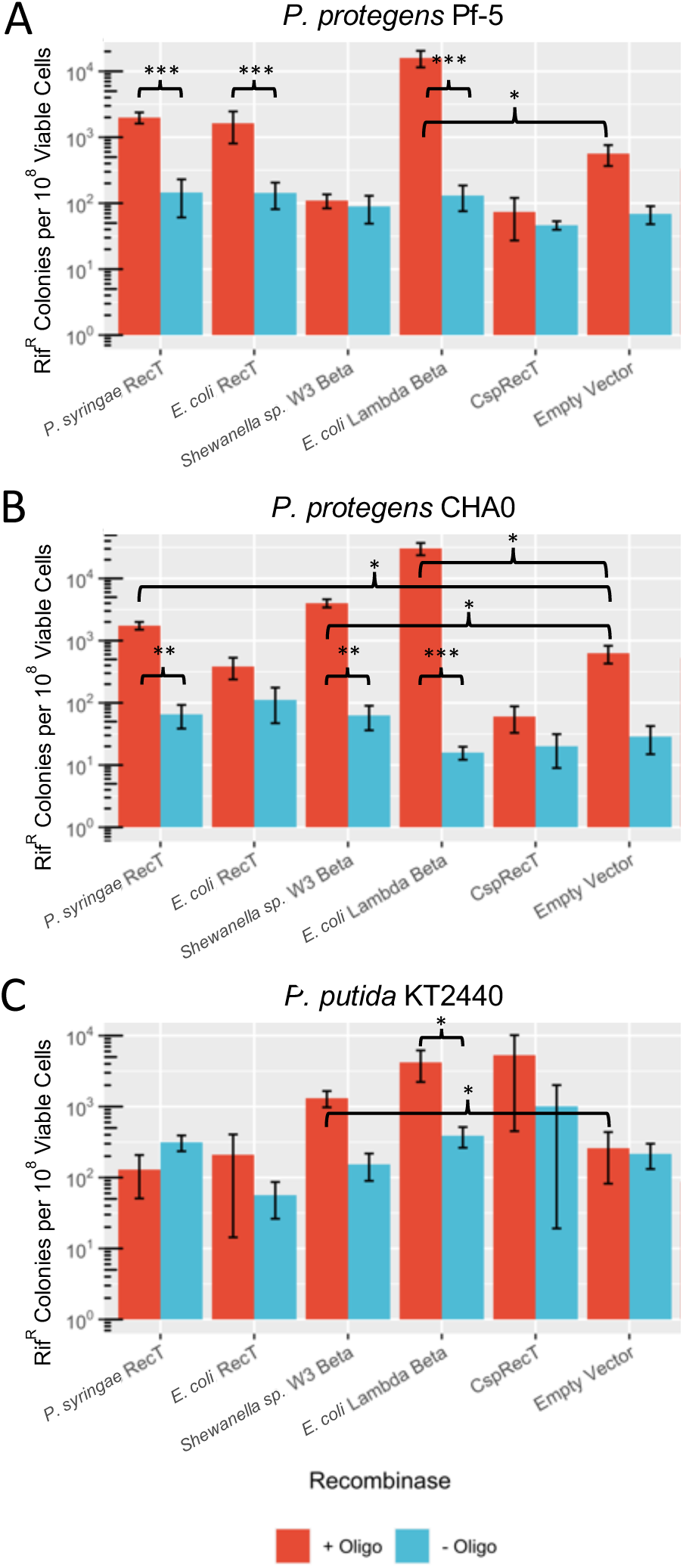
Comparison of SSAPs across *Pseudomonas* spp. Log phase cultures of (A) *P. protegens* Pf-5, (B) *P. protegens* CHA0, and (C) *P. putida* KT2440 expressing five candidate SSAPs or empty vector (pBBR1-MCS2) were electroporated with 15 µg of oligonucleotide encoding a D521P point mutation in *rpoB,* and the cell mixture recovered for 3.5 hours in LB before plating on rifampicin. Rif^R^ colonies and total viable colonies were counted after 2 days of growth. Significance values are indicated for a Mann-Whitney U test between two groups, where * p < 0.05; **p < 0.01; ***p < 0.001; and; ns, not significant.

Notably, candidate recombinase efficiency profiles varied across all three strains. The highest levels of recombineering frequencies within two out of three strains were achieved in the presence of SSAP λ-Red Beta (*E. coli*), at 1.6 x 10^-4^ for *P. protegens* Pf-5, 3.0 x 10^-4^ for *P. protegens* CHA0 (Figure 2). The most efficient recombinase for *P. putida* KT2440 was CspRecT, at a frequency of 5.3 x 10^-5^, followed closely by λ-Red Beta (Figure 2). Though CspRecT functioned well in *P. putida* KT2440, recombineering efficiencies using this SSAP were poor in both *P. protegens* strains. The next most efficient recombinase tested in *P. protegens* CHA0 and *P. putida* KT2440 was the λ-Red Beta-like SSAP from *Shewanella sp.* W3-18-1; however, this SSAP in *P. protegens* Pf-5 did not appear to improve recombineering efficiency above wildtype levels. The remaining two recombinases: RecT (*P. syringae* pv. *syringae* B728a) and RecT (*E. coli* Rac prophage) did not appear to significantly improve recombineering efficiencies in *P. putida* KT2440. *E. coli* RecT also did not support recombineering efficiencies above wildtype levels in *P. protegens* CHA0, but the *P. syringae* RecT SSAP resulted in a recombineering frequency of 1.7 x 10^-5^ (Figure 2). These SSAPs functioned similarly in *P. protegens* Pf-5, with recombineering frequencies of 2 x 10^-5^ and 1.6 x 10^-5^ for *P. syringae* RecT and *E. coli* RecT, respectively (Figure 2).

### Choice of mutation affects recombineering efficiencies

To investigate the effect of length and location of mutations on recombineering efficiencies, we designed additional oligonucleotides to target the Q518 residue in *rpoB*. Figure 3A depicts the different designs of oligonucleotides tested in the three *Pseudomonas* strains expressing *E. coli* Lambda Beta SSAP. We chose oligonucleotides with different nucleotide mismatch pairs, as well as different overall numbers of nucleotides.

**Figure 3.**
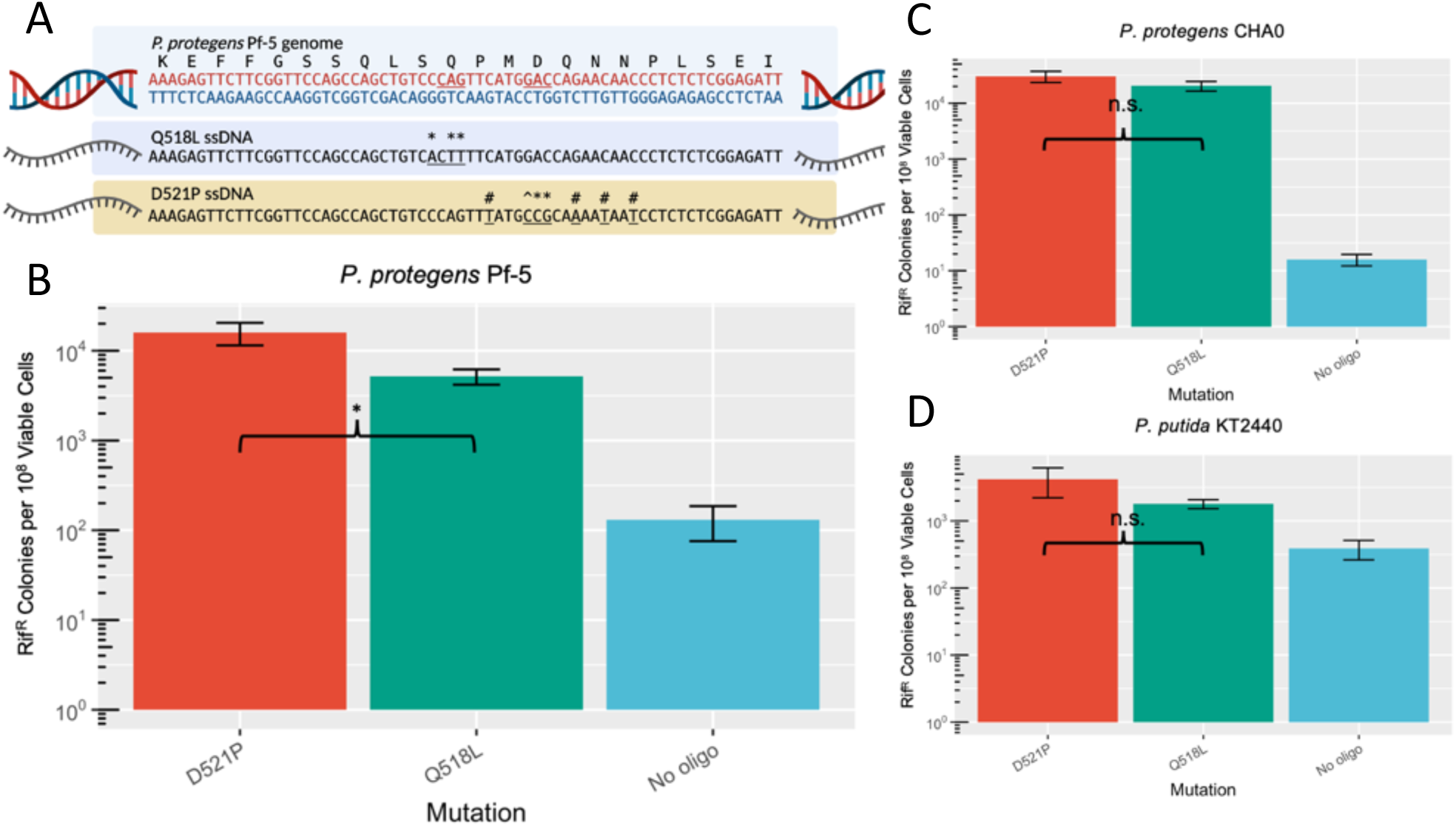
Comparison of *rpoB* point mutations across *Pseudomonas* spp. (A) ssDNA design of the Q518L and D521P point mutations. Single basepair mutations and codon changes are underlined. Individual basepair mutations are further denoted by *, #, and ^, where * indicates a transversion, # indicates a transition, and ^ indicates a rarely detected C:C mismatch. Log phase cultures of (B) *P. protegens* Pf-5, (C) *P. protegens* CHA0, and (D) *P. putida* KT2440 expressing *E. coli* λ Red Beta were electroporated with 15 µg of oligonucleotide encoding a D521P point mutation or Q518L in *rpoB.* Significance values are indicated for a Mann-Whitney U test between two groups, where * p < 0.05; **p < 0.01; ***p < 0.001; and; ns, not significant.

The addition of a few silent mutations flanking the D521P point mutation improved recombineering efficiencies in all three strains by a factor of ∼1.5, 2.3, or 3.0 for *P. protegens* CHA0, *P. putida* KT2440, and *P. protegens* Pf-5, respectively, compared to the Q518L mutation which is flanked by only one silent mutation (Figure 3). The degree to which additional silent mutations impacted recombineering efficiencies may reflect different levels of sensitivity to methyl-directed mismatch repair (MMR) in these strains. The two point mutations tested encode for individual basepair mismatches that may enable different levels of MMR evasion. The Q518L oligonucleotide encodes C → A, A → T, and G → T mutations, while the D521P oligonucleotide encodes C → T, G → A, and two C → T mutations as well as A → C, C → G and G → C mutations, the latter of which yields a C:C mismatch. In *E. coli*, C:C mismatches go nearly undetected by MMR machinery, which lead to an approximate 30-fold increase in subsequent recombineering efficiencies (16, 35). The C:C mismatch as well as increased mismatch basepairing in the D521P oligonucleotide may have diminished detection from the *Pseudomonas* MMR system, though the effects we see are not as improved as those reported in *E. coli*. *P. putida* MMR machinery has shown the least recognition for A:G and C:C mismatches (36), which also supports the higher recombineering rates for the D521P oligonucleotide observed in this study.

The Q518L mutation itself lends a contrasting phenotype in *P. protegens* strains between targeted recombineering mutants compared to spontaneous rifampicin-resistance mutants due to a growth defect (Supplementary Figure S3). In these experiments, visible colonies after two days of growth are the spontaneous Rif^R^ mutants, whereas smaller colonies visible after four days of growth are the Q518L mutants based on PCR screening (n=19). Growth assays comparing Q518L variants to wildtype and vector-carrying strains showed a two-fold increase in doubling time for both *P. protegens* strains CHA0 (68.6 ± 2.8 to 125.5 ± 7.7 minutes) and Pf-5 (67.1 ± 2.9 to 128.2 ± 2.8 minutes). The Q518L mutation did not result in a growth phenotype in *P. putida*.

### Recombineering efficiencies are influenced by oligonucleotide availability

A well-established phenomenon of recombineering is the improvement of recombinant frequency as oligonucleotide load increases until a saturating concentration is reached (4, 8, 34). We aimed to determine the saturation levels of oligonucleotide within these *Pseudomonas* strains to further optimize recombineering efficiencies. For all three strains, an oligonucleotide load of 15 μg is saturating, at an approximate copy number of 3 x 10^5^ per cell (Figure 4).

**Figure 4.**
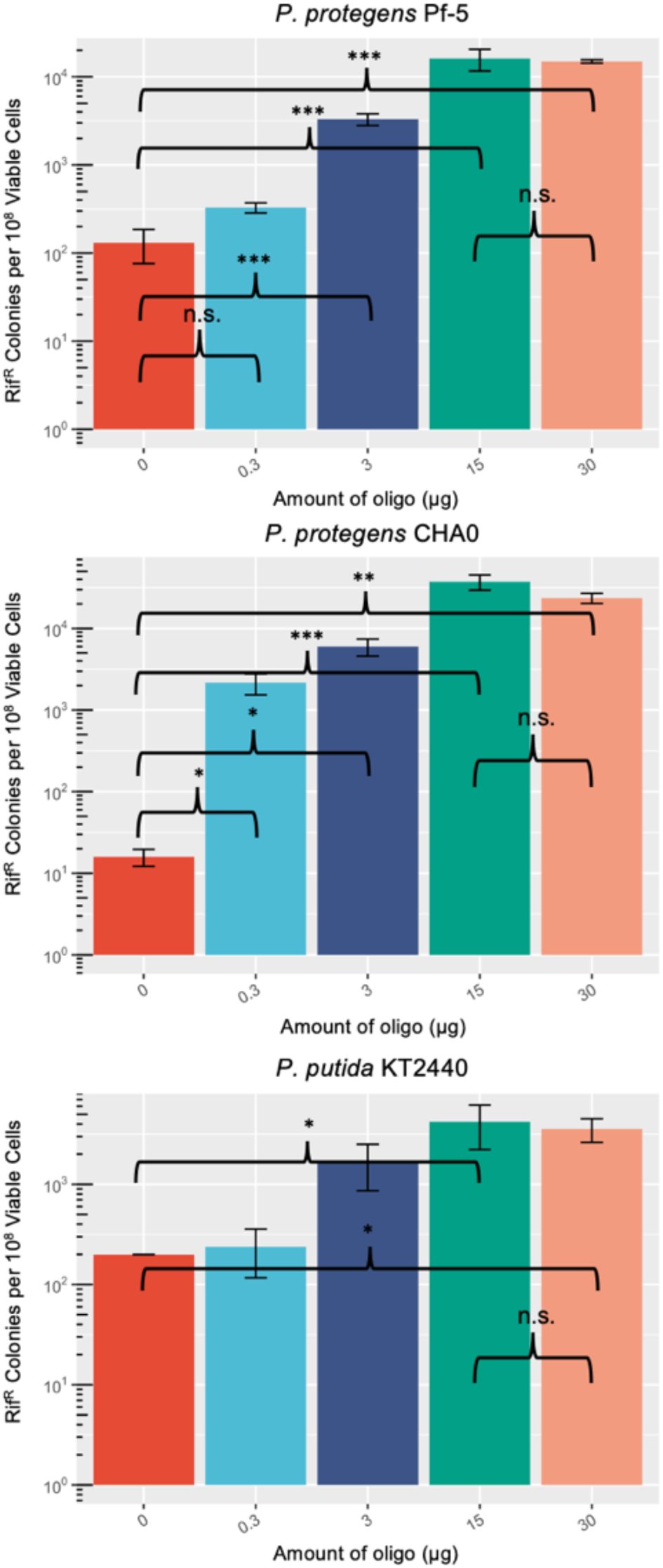
Effect of ssDNA amount on recombineering efficiency. Log phase cultures of *P. protegens* Pf-5, *P. protegens* CHA0, and *P. putida* KT2440 expressing *E. coli* λ Red Beta were electroporated with 0, 0.3, 3, 15, or 30 µg of oligonucleotide encoding a D521P point mutation in *rpoB,* and the cell mixture recovered for 3.5 hours in LB before plating on rifampicin. Rif^R^ colonies and total viable colonies were counted after 2 days of growth. Significance values are indicated for a Mann-Whitney U test between two groups, where * p < 0.05; **p < 0.01; ***p < 0.001; and; ns, not significant.

Many ssDNA recombineering protocols call for the addition of phosphorothioate bonds on the ends of recombineering oligonucleotides to prevent exonuclease degradation, which would theoretically increase the half-life of oligonucleotide within the cell. Typically, four phosphorothioate bonds are designed at the 5’ end of the recombinogenic oligonucleotide (2, 4, 17, 30). We tested the effect of phosphorothiolation using solely the *P. protegens* strains as they exhibited higher levels of recombineering efficiencies that were promising for further optimization (Figures 2, 4). We observed 3.3-fold and 1.3-fold improvements in the average recombineering efficiency using 5’ only phosphorothiolation or both 5’ and 3’ phosphorothiolation in *P. protegens* CHA0 (Figure 5). In *P. protegens* Pf-5, phosphorothiolation of the 5’ end only improved recombineering efficiencies 2.5-fold, whereas 5’ and 3’ treatment reduced efficiencies by 0.6-fold (Figure 5). We did not observe statistically significant improvements in recombineering efficiencies that have been reported in other studies (2, 30); however, testing the effects of phosphorothiolation at lower oligonucleotide concentration levels may reveal a more dramatic effect.

**Figure 5.**
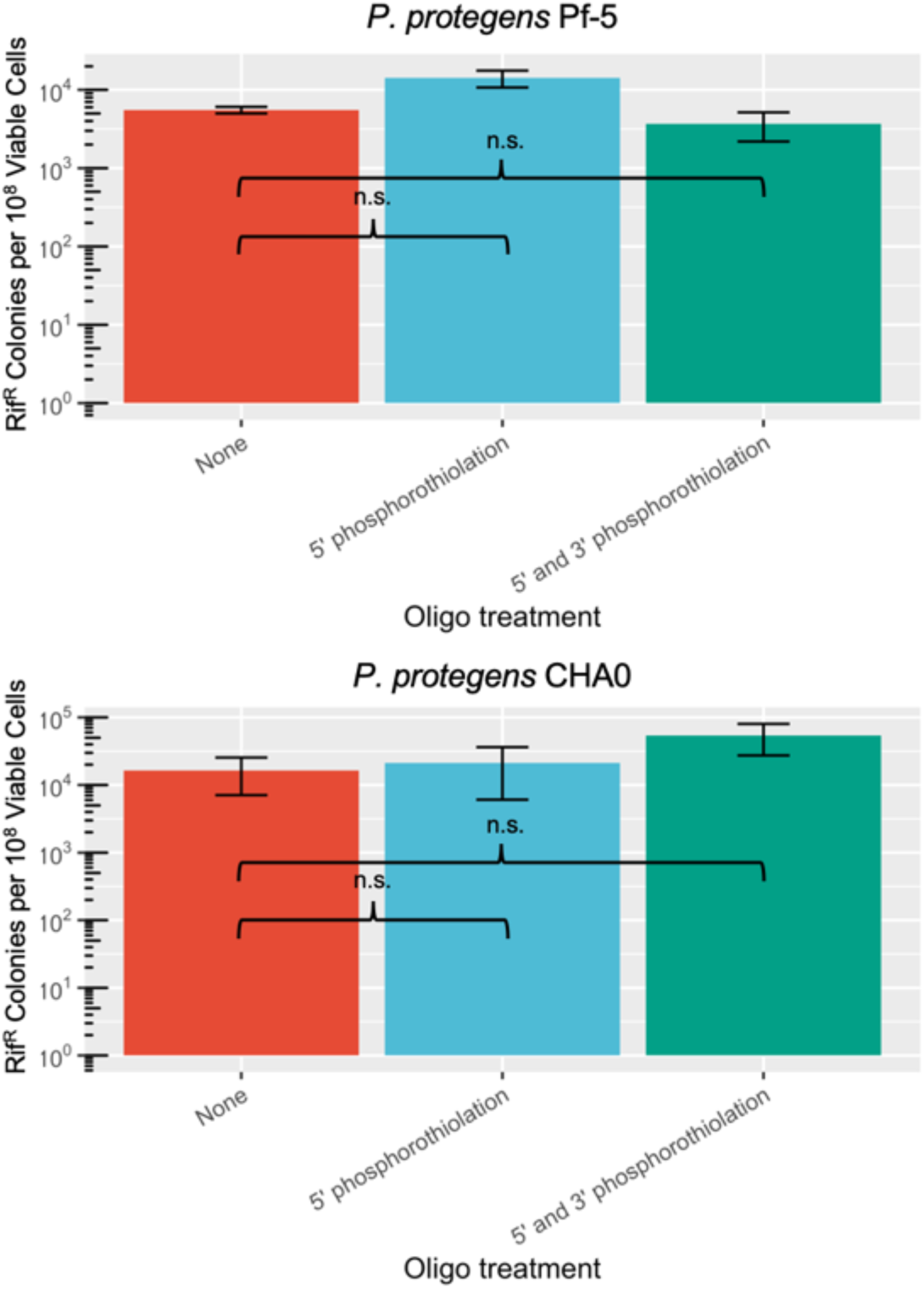
Effect of phosphorothiolation of oligonucleotide. Log phase cultures of *P. protegens* Pf-5 and *P. protegens* CHA0 expressing *E. coli* λ Red Beta were electroporated with 15 µg of oligonucleotide encoding a D521P point mutation without phosphorothioate bonds, with 4 5’ end phosphorothioate bonds, or 4 5’ and 3’ each phosphorothioate bonds, and the cell mixture recovered for 3.5 hours in LB before plating on rifampicin. Rif^R^ colonies and total viable colonies were counted after 2 days of growth. Significance values are indicated for a Mann-Whitney U test between two groups, where * p < 0.05; **p < 0.01; ***p < 0.001; and; ns, not significant.

## Discussion

Recombineering with SSAPs is a relatively easy and rapid method to introduce targeted mutations into a host genome. The flexibility of this genome engineering method has made it widely attractive to implement systems across several genera (11, 30, 38–41). In adapting a recombineering system to a new strain background, several recombineering bottlenecks should be addressed to improve efficiencies: most notably improving recombinogenic DNA availability and determining efficient SSAPs. Here we investigated methods to improve recombineering efficiencies in three *Pseudomonas* strains through manipulation of these common recombineering bottlenecks and demonstrated that strains within the same genera respond very differently to identical recombineering platforms.

A critical step in the recombineering pipeline is the efficient uptake of ssDNA. We found that transformation competencies of these strains do vary, impacting the achievable recombineering efficiencies. Using the electroporation procedure developed in this study, the corresponding transformation efficiencies for the *P. protegens* strains were similar, at approximately 5 x 10^8^ transformants per µg plasmid DNA (Figure S1). The transformation efficiency for *P. putida* KT2440, however, was several orders of magnitude lower at 2 x 10^6^ transformants per µg DNA. This discrepancy alone can account for the diminished recombineering efficiency we observed in *P. putida* KT2440. While we did examine the effect of electroporation buffer on transformation efficiency (Figure S1), several other variables could be optimized to improve electrotransformation and thus recombineering efficiencies in these strains.

We examined the effects of two unique mutations in the *rpoB* gene and found that the addition of flanking silent mutations improved recombineering efficiencies across all three strains. According to studies performed in model strains of *E. coli*, MMR recognition is generally overcome by the introduction of a minimum of 4 mismatched bases (42, 43). Little is known about the MMR constraints in *Pseudomonas*, though the results we see in this study do support what has been reported in *P. putida* EM42 (37). We also investigated the effects of oligonucleotide availability via substrate load amount and phosphorothiolation. In all strains, the dose-dependent trend of recombineering efficiency reached saturation at 15 µg of ssDNA. Phosphorothiolation of solely the 5’ end of the oligonucleotide appeared to slightly improve recombineering efficiencies; however, this variable did not seem as impactful as what has been reported in other organisms, which is consistent with a recent finding in the *P. putida* EM42 strain (44). This may indicate lower levels of endogenous exonucleases in these strains, or that at high concentrations (15 µg of ssDNA) the effect of oligo protection is less impactful.

The choice of SSAP remained the single most important factor affecting recombineering efficiency, resulting in frequencies of recombineering across several orders of magnitude (Figure 2). In this study we tested five recombinases that had either been shown to function in these or other strains of *Pseudomonas*, or other related environmental species like *Shewanella* (7, 12, 26, 27, 34). Interestingly, we found that the most efficient SSAPs were different between the strains: λ-Red Beta for *P. protegens* Pf-5 and CHA0, and CspRecT for *P. putida* KT2440. While it is known that strain to strain variability affects the successful portability of genetics systems, this is further evidence of the difficulty to widely adapt a recombineering system across strains even of the same genus. Additionally, the poor performance of CspRecT in the *P. protegens* strains is surprising, as CspRecT was reported to far outperform *E. coli* λ-Red Beta in both *E. coli* and *P. aeruginosa* backgrounds.

Recent studies have discussed a potential link affecting recombineering efficiencies between a candidate SSAP and the host’s single-stranded DNA binding protein (SSB) (11). It was found that specific combinations of SSBs and SSAPs resulted in improved recombineering efficiencies that were linked to the seven C-terminal amino acid sequence of the SSB. To investigate whether this interaction may have influenced the recombineering efficiencies we saw in this study, we performed an alignment of SSBs from several *Pseudomonas* strains (Figure S4). Notably, we found that all three strains investigated in this study shared an identical seven C-terminal amino acid sequence. Additionally, the SSBs from *P. protegens* Pf-5 and *P. protegens* CHA0 shared 100% similarity, yet the recombinase efficiency profiles between these two strains had significant differences, specifically when using the W3 Beta SSAP from *S. sp.* W3-18-1. While the relationships between SSAPs and SSBs may provide guidance on which SSAPs may be more successful in a particular host, the strain-to-strain variation we see in this study indicate that additional effects likely still influence the success of a candidate recombinase.

The results presented here emphasize the value of multiple strain studies, especially in the context of building gene editing technologies in non-model organisms. We found that recombineering efficiencies varied widely from a selection of *Pseudomonas spp.* under identical conditions including the expressed SSAP and mutation design. In the context of developing strains for potential uses as commercial genetically engineered organisms, initial manipulation studies that investigate several relevant strains is incredibly valuable. Such studies would be valuable for both identifying more responsive candidates as well as developing methodologies that function well across a genus or even more distant organisms. Given the many unknowns associated with research in non-model systems, this approach is a useful framework for developing recombineering methods when there is limited information about the molecular genetics of the host.

## Acknowledgements

This work was supported by the US Department of Energy, Office of Biological and Environmental Research, Genomic Science Program Lawrence Livermore National Laboratory’s Secure Biosystems Design Scientific Focus Area under grant award no. SCW1710.

